## Supplementary data for "An Arginine-Rich Motif in the ORF2 Capsid Protein Regulates the Hepatitis E Virus Lifecycle and Interactions with the Host Cell"

### Supporting Figures

**S1 Fig. Colocalization analysis of ORF2 with the Golgi marker 130 (GM130) in PLC3/HEV-p6 cells expressing ORF2wt or ARM/SP mutants.** PLC3/HEV-p6 cells expressing ORF2wt or ARM/SP mutants were analyzed by indirect immunofluorescence at 18 h.p.e. Cells were analyzed by confocal microscopy (magnification x63). Representative images are shown. Scale bar, 20 µm. Pearson's correlation coefficients were calculated using JACoP plugin from ImageJ software using the whole cell as ROI (mean ± S.D., $n \geq 30$ cells, Kruskal-Wallis with Conover's test). ***$p < 0.001$, ****$p < 0.0001$. Red = ORF2; Green = GM130; Blue = DAPI.

**S2 Fig. Colocalization analysis of ORF2 with the Importin-α1 in PLC3/HEV-p6 cells expressing ORF2wt and ARM/SP mutants.** PLC3/HEV-p6 cells expressing ORF2wt or ARM/SP mutants were analyzed by indirect immunofluorescence at 18 h.p.e. Cells were analyzed by confocal microscopy (magnification x63). Representative images are shown. Scale bar, 20 µm. Pearson's correlation coefficients were calculated using JACoP plugin from ImageJ software using the whole cell as ROI (mean ± S.D., $n \geq 30$ cells, Kruskal-Wallis with Conover's test). ****$p < 0.0001$. Red = ORF2; Green = Importin-α1; Blue = DAPI.

**S3 Fig. Transcriptomic analysis.** (A) Representative images of ORF2 subcellular localization in PLC3/HEV-p6 cells expressing ORF2wt, 5R/5A mutant, ΔORF3 mutant or Mock-electroporated PLC3 cells at 18 h.p.e, the time point at which RNA extraction and transcriptomic analyses were performed. Cells were analyzed by confocal microscopy (magnification x63). Red = ORF2; Blue = DAPI. (B) Representative images showing the ORF3 expression in cells used for transcriptomic analyses. Since ORF3 is poorly expressed at early time points post-electroporation, electroporated cells were fixed at 6 d.p.e. and processed for ORF2 and ORF3 staining. Cells were analyzed by confocal microscopy (magnification x63). Red = ORF2; Green = ORF3; Blue = DAPI. Scale bar, 20μm. (C) Intracellular RNAs were quantified at 18 h.p.e by RT-qPCR using CCL2, CCL20, CXCL1, CXCL2, NFKBIA, TNFAIP2 or TNFAIP3-targeting probes. Intracellular HEV RNA levels were controlled using an ORF1-targeting probe. $n \geq 6$, Kruskal-Wallis with Dunn's test for CCL2 and TNFAIP2 and ANOVA with Dunnett's test for the other genes. *$p < 0.05$, **$p < 0.01$, ****$p < 0.0001$.

**S4 Fig. Colocalization analysis of ORF2 with the exportin CRM1 in PLC3/HEV-p6 cells expressing ORF2wt or NES mutants.** PLC3/HEV-p6 cells expressing ORF2wt or NES mutants were analyzed by indirect immunofluorescence at 48 h.p.e. Cells were analyzed by confocal microscopy (magnification x63). Representative images are shown. Scale bar, 20 μm. Red = ORF2; Green = CRM1; Blue = DAPI. Pearson's correlation coefficients were calculated using JACoP plugin from ImageJ software using the whole cell as ROI (mean ± S.D., $n \geq 30$ cells, Kruskal-Wallis with Conover's test). ***$p < 0.001$, ****$p < 0.0001$.

**S5 Fig. Addressing of C2 constructs.** Schematic representation of the ORF2wt protein. SP ORF2 residues are shown in blue. ARM residues are highlighted in red. (A) H7-T7-IZ cells were transfected with pTM plasmids expressing ORF2wt or C2 constructs. In these constructs, the first half of the ORF2 SP (SP1) was deleted. Twenty-four hours post-transfection, cells were fixed and processed for ORF2 staining (in red). Nuclei are in blue. Representative confocal images are shown together with

ORF2/DAPI merge images. Blue dots observed in some pictures are DAPI-stained transfected plasmids. A schematic representation of each construct is shown on the left. Scale bar, 20$\mu$m. (B) Nuclear-to-cytoplasmic ORF2 staining ratio in H7-T7-IZ cells expressing ORF2wt or C2 constructs. Quantification was done using ImageJ software (mean $\pm$ S.D., $n \geq 30$ cells, Kruskal-Wallis with Conover's test). $**p < 0.01$, $****p < 0.0001$. (C) Subcellular fractionation of H7-T7-IZ cells expressing ORF2wt, C2 constructs or the pTM empty vector, at 24h post-transfection. Fractionation was done using the subcellular protein fractionation kit for cultured cells. ORF2 proteins were detected by WB with the 1E6 Ab. Tubulin, Calnexin (CNX) and Lamin B1 were also detected to control the quality of fractionation. Molecular mass markers are indicated on the right (kDa).

**S6 Fig. Dose-response curves of PLC3 cells treated with the different drugs used in this study.** Cell viability was determined by a MTS based assay. Cells treated with the solvent (DMSO or Ethanol) were used as a control and set to 100%. For LepB and Verd, blue arrows indicate the concentration used for subsequent experiments. Times of treatment are indicated.

### Supporting Materials and Methods

**RNA extraction and quantification.** HEV RNA levels were quantified by RT-qPCR using primers (5’-GGTGGTTTCTGGGGTGAC-3’ (F) and 5’-AGGGGTTGGTTGGATGAA-3’ (R)) and a probe (5’-FAM-TGATTCTCAGCCCTTCGC-TAMRA-3’) that target a conserved 70 bp region in the ORF2/3 overlap.

In **S3 Fig,** HEV RNA levels were quantified by RT-qPCR using primers (5’-AAGACATTCTGCGCTTTGTT-3’ (F) and 5’- TGACTCCTCATAAGCATCGC-3’ (R)) and a probe (5’-FAM- CCGTGGTTCCGTGCCATTGA-TAMRA-3’) that target a conserved region of ORF1. HEV RNAs were extracted from culture supernatants with the QIAmp viral RNA mini kit (Qiagen) and from cells with the Nucleospin RNA Plus kit (Macherey & Nagel). Retrotranscription was performed using the AffinityScript Multiple temperature cDNA synthesis Kit (Agilent Technologies) according to manufacturer’s instructions. Amplifications were done with a Quant Studio 3 apparatus (Applied Biosystems) and Taqman universal master mix no AmpErase UNA (Applied Biosystems). Cellular gene RNA levels were quantified by RT-qPCR using in-home primers (see **S2 Table**) and standards. Total cellular RNAs were extracted using TRIzol (Invitrogen) according manufacturer’s instructions and processed for retrotranscription using the High-capacity reverse transcription kit (Applied Biosystems). Amplifications were done with a Quant Studio 3 apparatus (Applied Biosystems) and SYBRGreen PCR Master Mix (Applied Biosystems).

**Transcriptomic analysis.** PLC3 cells were electroporated with HEV-p6-wt, HEV-p6-5R/5A, HEV-p6-ΔORF3 RNAs or no RNA (mock). At 18 h.p.e. total cellular RNAs were extracted using TRIzol (Invitrogen) according manufacturer's instructions. RNA integrity and purity were verified using the Agilent Bioanalyzer system (Agilent Technology). Two µg of total RNA were treated with 2 units of DNaseI (Sigma Aldrich) during 10 min before purification on Nucleomag NGS cleanup beads (Macherey Nagel). Oligonucleotide microarrays for human whole genome (G4858A design 072363, 8x60k chips SurePrint G3 unrestricted GE, Agilent Technologies) were used for global gene expression analysis. Two hundred ng of total RNA was used in the Agilent Quick-Amp Labeling kit according to manufacturer's instructions. After purification using an RNeasy Mini Kit (Qiagen), cRNA yield and incorporation efficiency (specific activity) into the cRNA were determined using a NanoDrop 2000 (Thermo Scientific) spectrophotometer. For each sample, a total of 600 ng of cRNA was fragmented and hybridized overnight at 65°C. After hybridization, slides were washed before being scanned on a SureScan Microarray Scanner (Agilent Technologies) and further processed using Feature Extraction v10.7.3.1 software. The resulting text files were uploaded into language R v4.0.3 and analyzed using the LIMMA package (Linear Model for Microarray Data) [1,2]. A 'within-array' normalization was performed using LOWESS (locally weighted linear regression) to correct for dye and spatial effects [3]. Moderate *t*-statistic with empirical Bayes shrinkage of the standard errors [4] was then used to determine significantly modulated genes. Statistics were corrected for multiple testing using a false-discovery rate approach. Protein-protein interactions network was generated using STRING database [5]. Gene ontology enrichment was performed using Metascape resource [6] (www.metascape.org) on the significantly modulated genes to identify pathways significantly modulated by either wild-type or mutants.

### Supporting tables

**S1 Table: Primers of use to generate ORF2/CD4 chimeras/mutants.**

| Name (Orientation) | Sequence | Corresponding mutant/group |
|---|---|---|
| HEV-1 (Fw) | TTTCTGCCTATGCTGCCCGCGCCACCGGCCGGCCAGCCGTCTGGCGCTGCTGCTGGGCGGCGCAGCGGCGGTG CCGGCGGTGGTTTCTGGGGTGACAGG | 3R/3A |
| HEV-2 (Rev) | CCTGTCACCCCAGAAACCACCGCCGGCACCGCCGCTGCGCCGCCCAGCAGCAGCGCCAGACGGCTGGCCGGCC GGTGGCGCGGGCAGCATAGGCAGAAA | |
| HEV-3 (Fw) | CGTCGTCGTGGGGCGGCCAGCGGCGGTGC | 2R/2A |
| HEV-4 (Rev) | GCACCGCCGCTGGCCGCCCCACGACGACG | |
| HEV-5 (Fw) | CCGTCTGGCGCTGCTGCTGGGGCGGCCAGCGGCGGTGCCGGCGGTGCACCGCCGCTGGCCGCCCCACGACGA CG | 5R/5A |
| HEV-6 (Rev) | ACCGCCGGCACCGCCGCTGGCCGCCCCAGCAGCAGCGCCAGACGG | |
| HEV-7 (Fw) | TCTGGCCGTCGTCGTGCGCGGCGCAGCGGCGGT | G/A |
| HEV-8 (Rev) | ACCGCCGCTGCGCCGCGCACGACGACGGCCAGA | |
| HEV-9 (Fw) | CCCGCGCCACCGGCCGGCCAGCGGCGTCGCCGTCGTCGTGGGCGGCGC | PSG/3R |
| HEV-10 (Rev) | GCGCCGCCCACGACGACGGCGACGCCGCTGGCCGGCCGGTGGCGCGGG | |
| HEV-11 (Fw) | CAGCCGTCTGGCCGTCGTCG | ΔSP |
| HEV-12 (Rev) | ACGGCCAGACGGCTGCATGGTGATCCCATGGGCGATGCAACA | |
| HEV-13 (Fw) | CTGCCTATGCTGCCCGCGCCACCGGCCGGCCAG | ΔSP1 |
| HEV-14 (Rev) | GGGCAGCATAGGCAGCATGGTGATCCCATGGGCGATGCAACAAACATGTTATTCATT | |
| HEV-15 (Fw) | CCTGCCCCCTCACGCCCTTTCTCAGTCGCTCGCGCTAACGATGCTTTGTGGGCCTCCGCCACTGCCGCTGAGTAC GATCAGGCTACG | NES9 |
| HEV-16 (Rev) | CGTAGCCTGATCGTACTCAGCGGCAGTGGCGGAGGCCCACAAAGCATCGTTAGCGCGAGCGACTGAGAAAGGGC GTGAGGGGGCAGG | |
| HEV-17 (Fw) | CAGCAGTATTCTAAGACATTTTATGTTGCCCCGGCCCGCGGGAAGGCGTCCGCTTGGGAGGCTGGCACAACTAG GGCCGGC | NES10 |
| HEV-18 (Rev) | GCCGGCCCTAGTTGTGCCAGCCTCCCAAGCGGACGCCTTCCCGCGGGCCGGGGCAACATAAAATGTCTTAGAAT ACTGCTG | |
| HEV-19 (Fw) | CGTACCCTAGGTTTGCAGGGTTGTGCAGCCCAGTCCACTGCTGCTGAGGCTCAGCGCGCTAAAACGGAGGTAGG CAAAACCCGGGAG | NES12 |
| HEV-20 (Rev) | CTCCCGGGTTTTGCCTACCTCCGTTTTAGCGCGCTGAGCCTCAGCAGCAGTGGACTGGGCTGCACAACCCTGCAA ACCTAGGGTACG | |
| HEV-21 (Fw) | GATATGTACAACCATGCTGCCTATGCTGCCCGCGCC | ORF2wt |
| HEV-22 (Fw) | GATATGTACAACCATGTGCCCTAGGGTTGTTCTGCTGCTGTTCTTCGTGTTTCTGCCTATGCTGCCCGCGCCACCG GCCGGCAGCGGCGGTGCCGGCGGTGGTTTCTGGGGTG | |
| HEV-23 (Fw) | GATA TGTACA ACC ATGTGCCCTAGGGTTGTTCTGC | C2 |

| | | |
|---|---|---|
| HEV-24 (Fw) | GATATGTACAACCATGCTGCCTATGCTGCCCGCGCCACCGGCCGGCAGCGGCGGTGCCGGCGGTGGTTTCTGGGG | |
| HEV-29(Fw) | GATATGTACAACCATGAACCGGGGAGTCCCTTTTAGGCACTTGCTTCTGGTGCTGCAACTGGCGCTCCTCCCAGCAGCCACTCAGGGA | C1 |
| HEV-30 (Fw) | GATATGTACAACCATGAACCGGGGAGTCCCTTTTAGGCACTTGCTTCTGGTGCTGCAACTGGCGCTCCTCCCAGCAGCCACTCAGGGACAGCCGTCTGGCCGTCGTCGTGGGCGGCGC | |
| HEV-31 (Fw) | GATATGTACAACCATGAACCGGGGAGTCCCTTTTAGGCACTTGCTTCTGGTGCTGCAACTGGCGCTCCTCCCAGCAGCCACTCAGGGACAGCCGTCTGGCGCTGCTGCTGGGGCGGCC | |
| HEV-32 (Fw) | GATATGTACAACCATGAACCGGGGAGTCCCTTTTAGGCACTTGCTTCTGGTGCTGCAACTGGCGCTCCTCCCAGCAGCCACTCAGGGACAGCGGCGTCGCCGTCGTCGTGGGCGGCGC | |
| HEV-33 (Fw) | GATAGAATTCACCATGTGCCCTAGGGTTGTTCTGCTGCTGTTCTTCGTGTTTCTGCCTATGCTGCCCGCGCCACCGGCCGGCAAGAAAGTGGTGCTGGGCAAAAAAGGGGATACAGTGG | C4 |
| HEV-34 (Fw) | ATAGAATTCACCATGTGCCCTAGGGTTGTTCTGCTGCTGTTCTTCGTGTTTCTGCCTATGCTGCCCGCGCCACCGGCCGGCCAGCCGTCTGGCCGTCGTCGTGGGCGGCGCAAGAAAGTGGTGCTGGGCAAAAAAGGG | |
| HEV-35 (Fw) | ATAGAATTCACCATGTGCCCTAGGGTTGTTCTGCTGCTGTTCTTCGTGTTTCTGCCTATGCTGCCCGCGCCACCGGCCGGCCAGCCGTCTGGCGCTGCTGCTGGGGCGGCCAAGAAAGTGGTGCTGGGCAAAAAAGGG | |
| HEV-36 (Fw) | ATAGAATTCACCATGTGCCCTAGGGTTGTTCTGCTGCTGTTCTTCGTGTTTCTGCCTATGCTGCCCGCGCCACCGGCCGGCCAGCGGCGTCGCCGTCGTCGTGGGCGGCGCAAGAAAGTGGTGCTGGGCAAAAAAGG | |
| HEV-37 (Fw) | GATAGAATTCACCATGAACCGGGGAGTCCCTTTTAGGCACTTGCTTCTGGTGCTGCAACTGGCGCTCCTCCCAGCAGCCACTCAGGGACAGCCGTCTGGCCGTCGTCGTGGGCGGCGC | C5 |
| HEV-38 (Fw) | GATAGAATTCACCATGAACCGGGGAGTCCCTTTTAGGCACTTGCTTCTGGTGCTGCAACTGGCGCTCCTCCCAGCAGCCACTCAGGGACAGCCGTCTGGCGCTGCTGCTGGGGCGGCC | |
| HEV-39 (Fw) | GATAGAATTCACCATGAACCGGGGAGTCCCTTTTAGGCACTTGCTTCTGGTGCTGCAACTGGCGCTCCTCCCAGCAGCCACTCAGGGACAGCGGCGTCGCCGTCGTCGTGGGCGGCGC | |
| HEV-42 (Rev) | GATAGGATCCTTAAGACTCCCGGGTTTTGCCTACC | ORF2 groups |
| HEV-43 (Rev) | GATAACGCGTTCATTAATTAGGCCTCTCGAGCTGC | CD4 groups |
| HEV-44 (Fw) | GCTGTTCTACTCTCGCCC | ORF2 sequencing |
| HEV-45 (Fw) | CGCTACCGCCCGCTGGT | |
| HEV-46 (Rev) | ACCAGCGGGCGGTAGCG | |
| HEV-47 (Rev) | GCATTCTCCACAGATGT | |
| HEV-48 (Rev) | ACTGTAAAGGCGAGTGGG | CD4 sequencing |
| HEV-49 (Fw) | TTCTGGGAAATCAGGGC | |

**S2 Table: Primers used for RT-qPCR of cellular genes.**

| Primer name / Target | Sequence |
|---|---|
| hCCL2Fw / CCL2 | CATGAAAGTCTCTGCCGCCC |
| hCCL2Rev / CCL2 | GGGCATTGATTGCATCTGGCTG |
| hCCL20Fw / CCL20 | TTGTGCGTCTCCTCAGTAAAAA |
| hCCL20Rev / CCL20 | TCCAACCCCAGCAAGGTTC |
| hCXCL1Fw / CXCL1 | CTTCCTCCTCCCTTCTGGTC |
| hCXCL1Rev/ CXCL1 | GAAAGCTTGCCTCAATCCTG |
| hCXCL2Fw / CXCL2 | GCTTCCTCCTTCCTTCTGGT |
| hCXCL2Rev / CXCL2 | GGGCAGAAAGCTTGTCTCAA |
| hNFKBIAFw / NFKBIA | AAGGCCAGGTCTCCCTTCAC |
| hNFKBIARev / NFKBIA | CAGCAGCTCACCGAGGAC |
| hTNFAIP2Fw / TNFAIP2 | CTACGCTGGCCGAGATCATT |
| hTNFAIP2Rev / TNFAIP2 | CTCAGGTGGCCTTTGCTGAA |
| hTNFAIP3Fw / TNFAIP3 | GGCCCGGAGAGGTGTTG |
| hTNFAIP3Rev / TNFAIP3 | TCTTCTGGAGTTCTCTCCCGT |

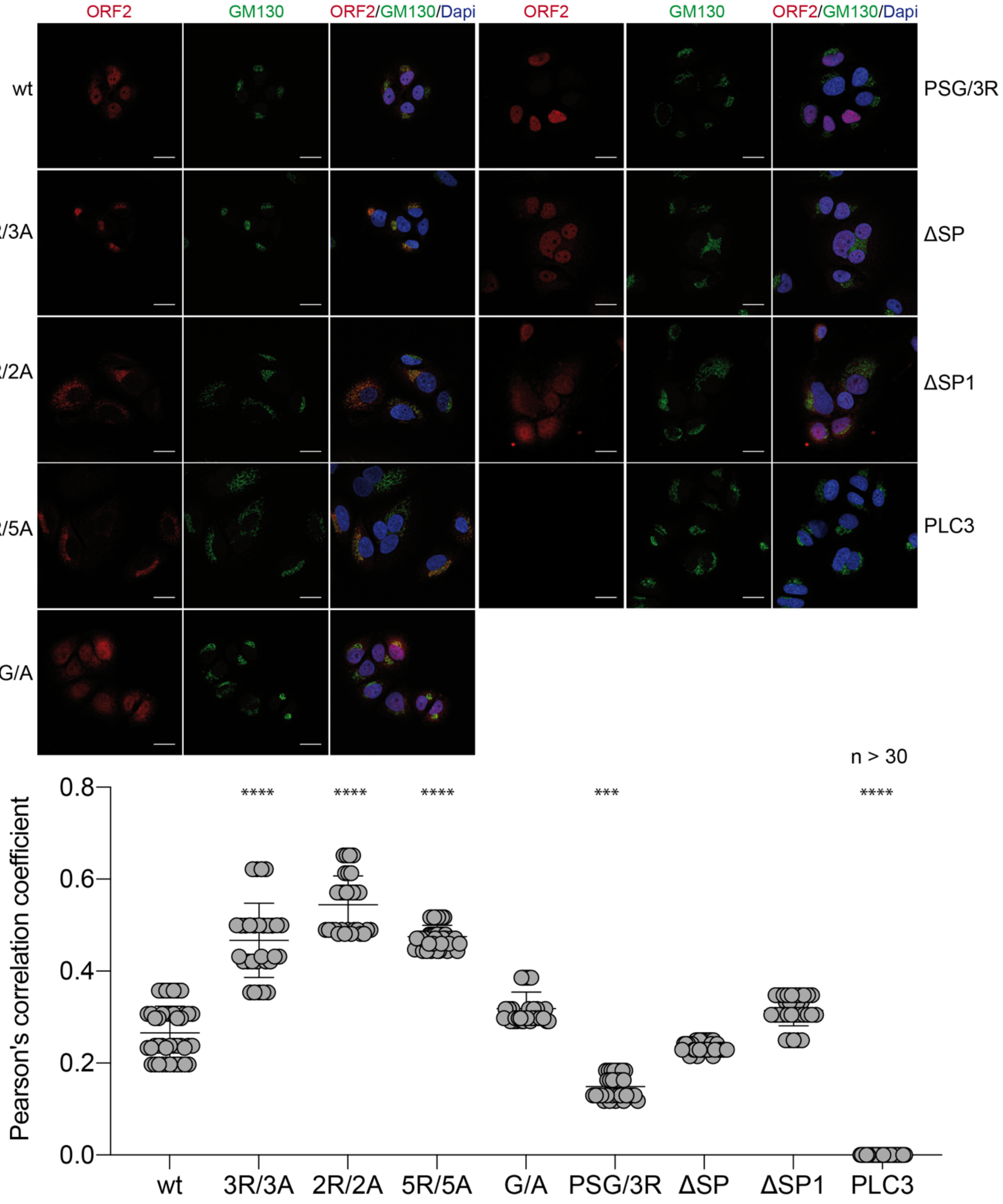
ORF2
GM130
ORF2/GM130/Dapi
wt
3R/3A
2R/2A
5R/5A
G/A
PSG/3R
ΔSP
ΔSP1
PLC3
n > 30
Pearson's correlation coefficient
0.8
0.6
0.4
0.2
0.0
****
***


**Fig S1**

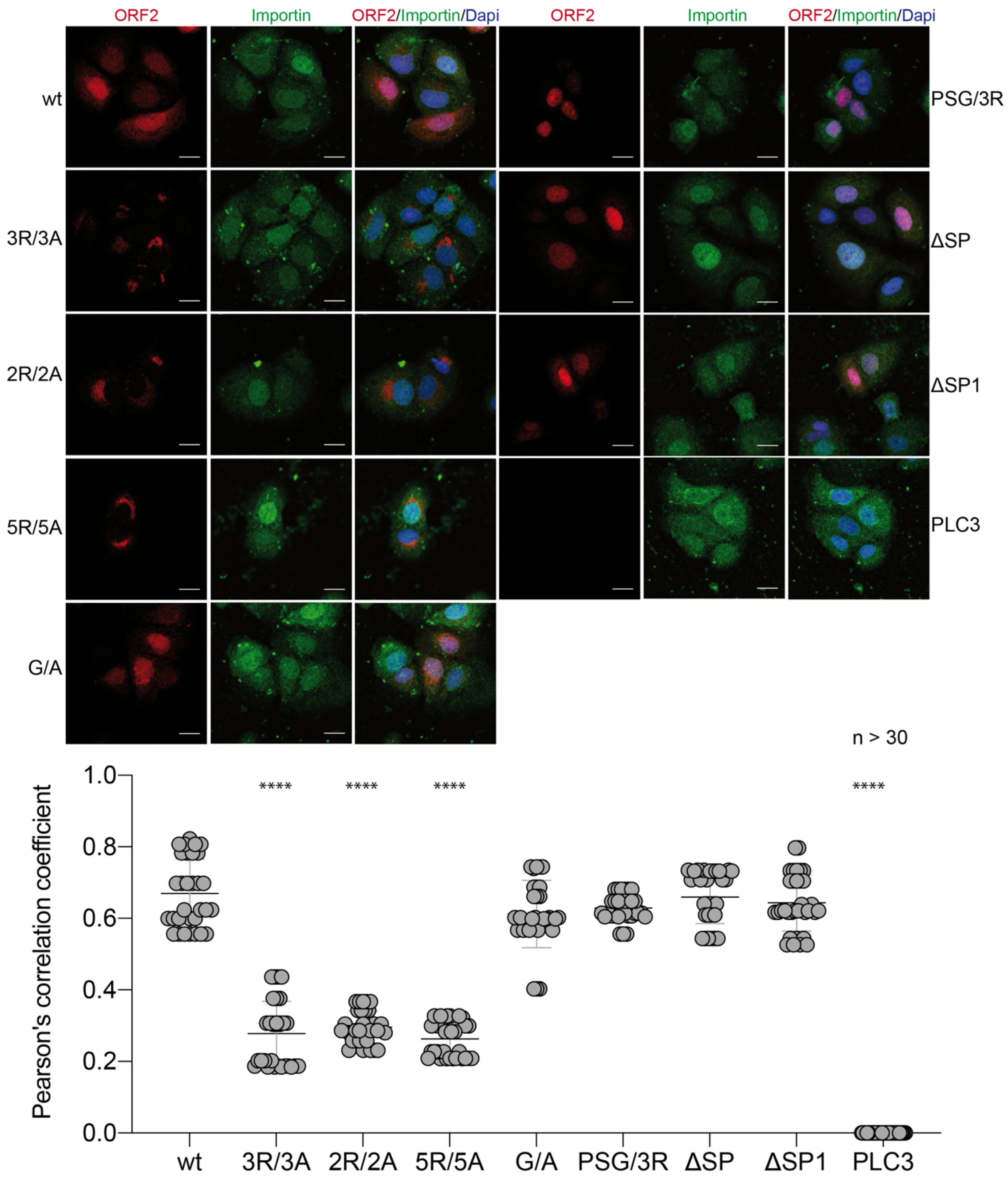
ORF2
Importin
ORF2/Importin/Dapi
ORF2
Importin
ORF2/Importin/Dapi
wt
3R/3A
2R/2A
5R/5A
G/A
PSG/3R
ΔSP
ΔSP1
PLC3
n > 30
Pearson's correlation coefficient
1.0
0.8
0.6
0.4
0.2
0.0
****
****
****
****
wt
3R/3A
2R/2A
5R/5A
G/A
PSG/3R
ΔSP
ΔSP1
PLC3

**Fig. S2**

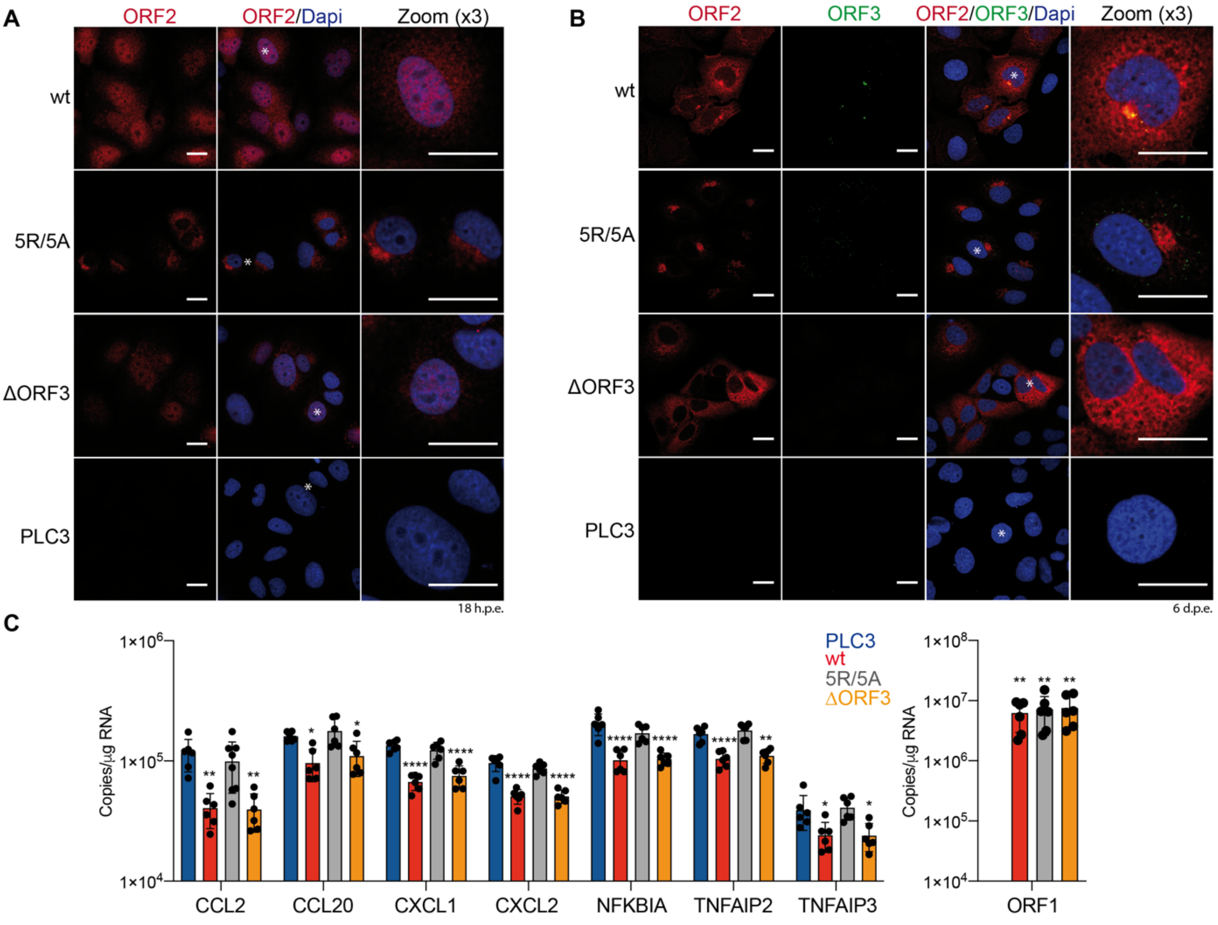
A
ORF2
ORF2/Dapi
Zoom (x3)
wt
5R/5A
ΔORF3
PLC3
18 h.p.e.
B
ORF2
ORF3
ORF2/ORF3/Dapi
Zoom (x3)
wt
5R/5A
ΔORF3
PLC3
6 d.p.e.
C
Copies/µg RNA
PLC3
wt
5R/5A
ΔORF3
CCL2
CCL20
CXCL1
CXCL2
NFKBIA
TNFAIP2
TNFAIP3
ORF1

**Fig. S3**

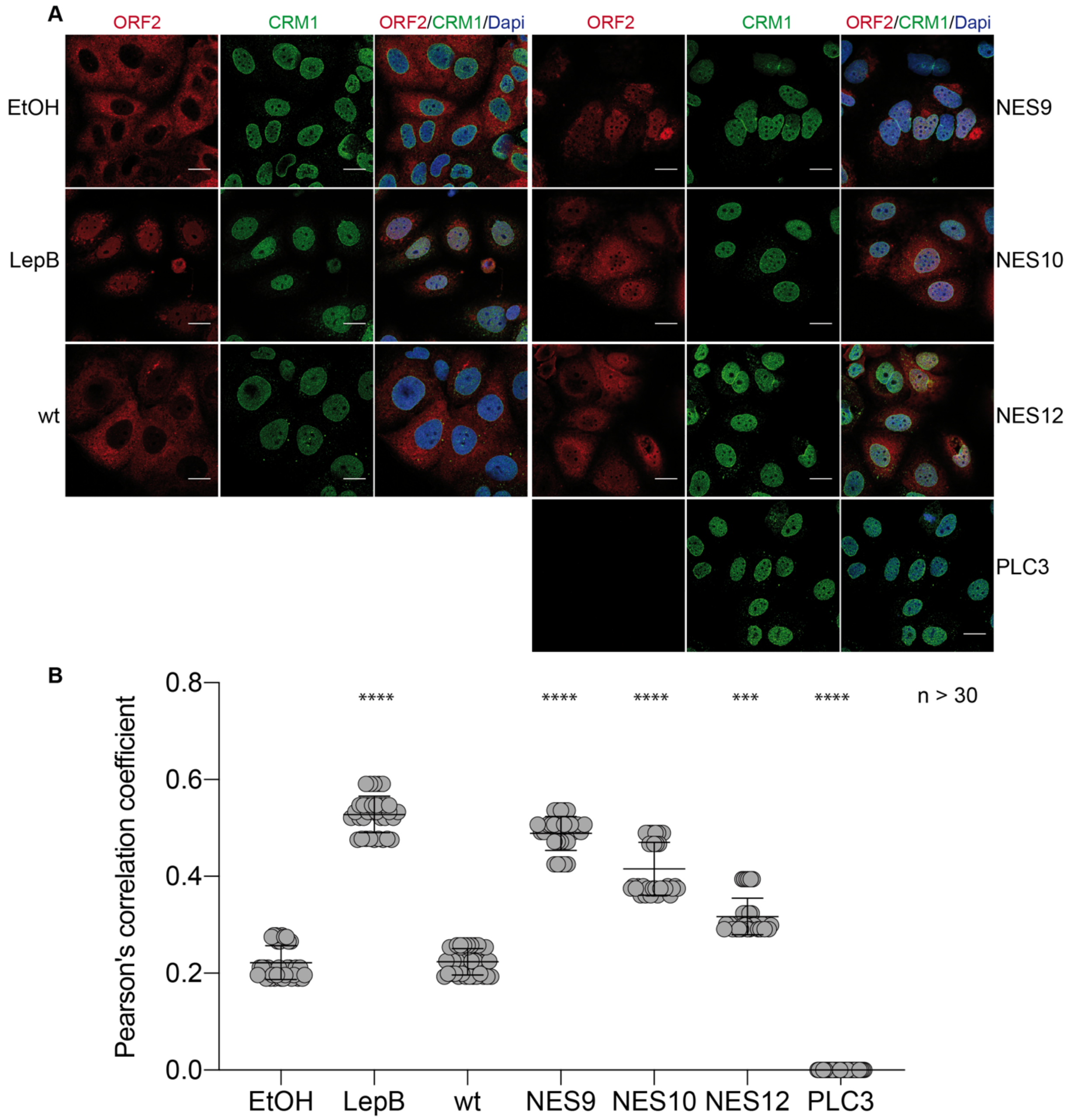
A
ORF2
CRM1
ORF2/CRM1/Dapi
EtOH
LepB
wt
NES9
NES10
NES12
PLC3
B
Pearson's correlation coefficient
0.0
0.2
0.4
0.6
0.8
****
****
****
***
****
n > 30
EtOH
LepB
wt
NES9
NES10
NES12
PLC3

**Fig. S4**

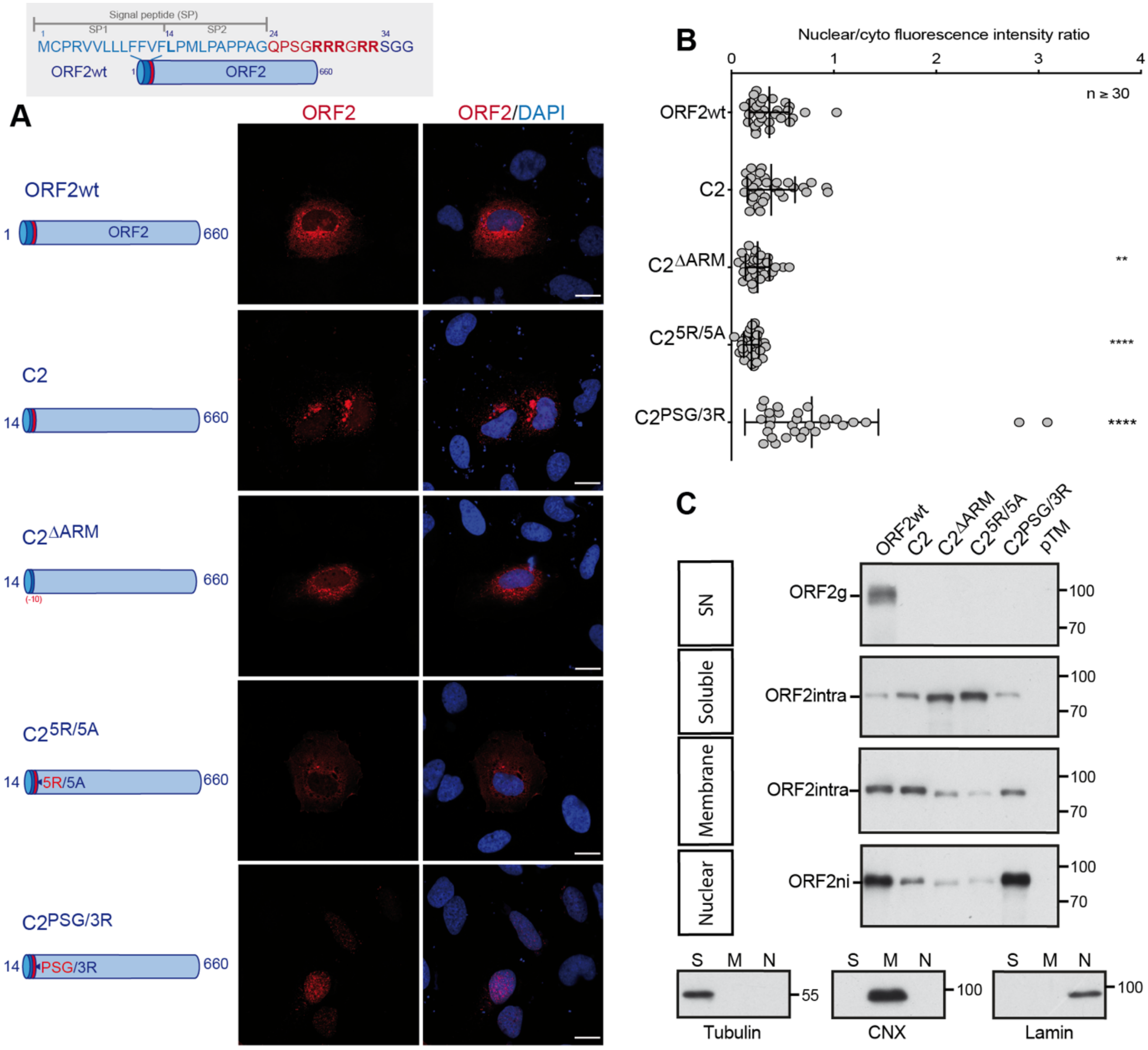
Signal peptide (SP)
SP1
SP2
MCPRVVLLLFFVFLPMLPAPPAGQPSGRRRGRRSGG
ORF2wt
ORF2
660
A
ORF2
ORF2/DAPI
ORF2wt
C2
C2ΔARM
C25R/5A
C2PSG/3R
5R/5A
PSG/3R
B
Nuclear/cyto fluorescence intensity ratio
n ≥ 30
**
****
****
C
ORF2g
ORF2intra
ORF2ni
SN
Soluble
Membrane
Nuclear
pTM
Tubulin
CNX
Lamin

**Fig. S5**

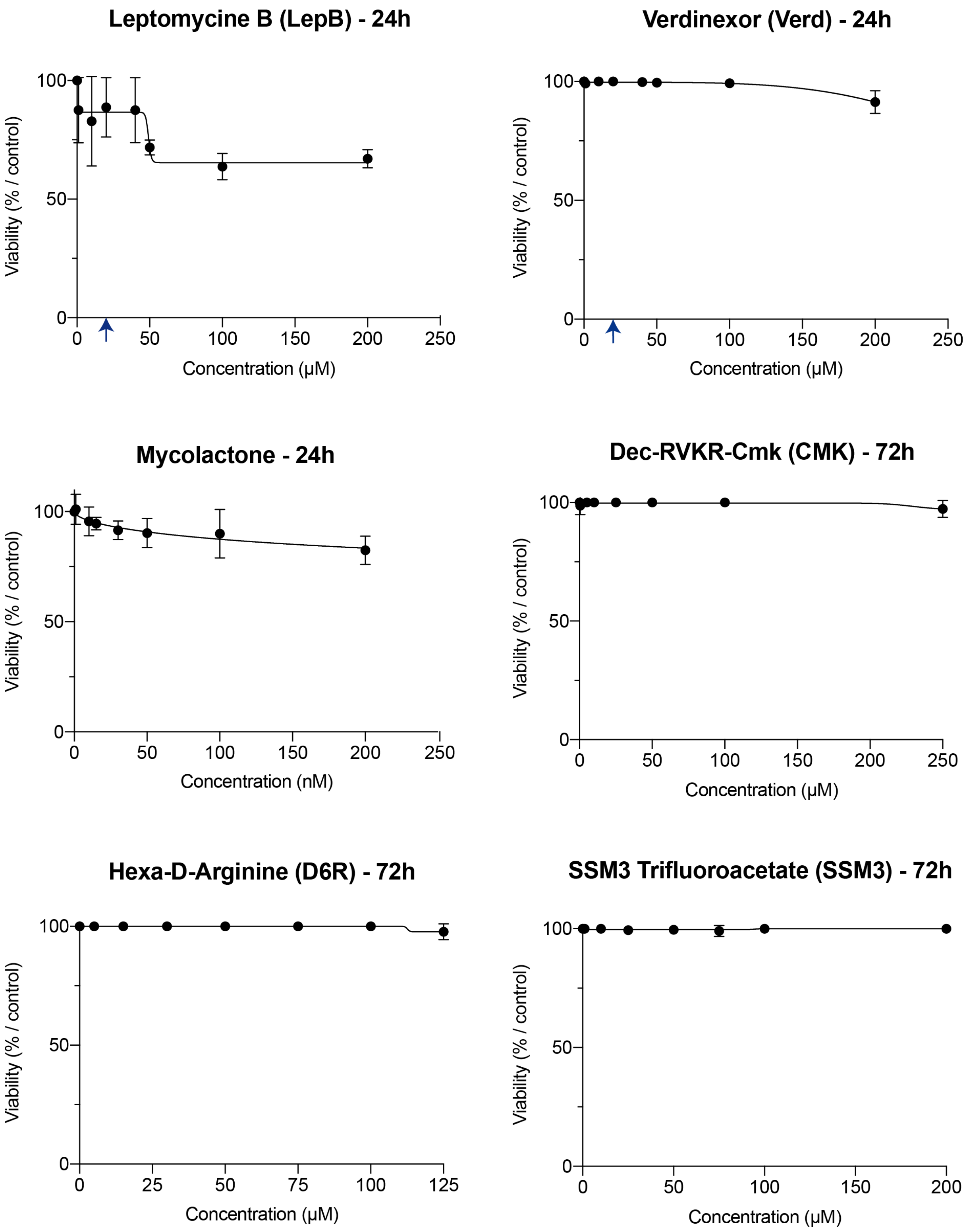

Leptomycine B (LepB) - 24h
Verdinexor (Verd) - 24h
Mycolactone - 24h
Dec-RVKR-Cmk (CMK) - 72h
Hexa-D-Arginine (D6R) - 72h
SSM3 Trifluoroacetate (SSM3) - 72h
Viability (% / control)
Concentration (µM)
Concentration (nM)
0
50
100
150
200
250
25
75
125


**Fig. S6**
